# Urinary biomarker and histopathological evaluation of vancomycin and piperacillin-tazobactam nephrotoxicity in comparison with vancomycin in a rat model and a confirmatory cellular model

**DOI:** 10.1101/568907

**Authors:** Gwendolyn M. Pais, Jiajun Liu, Sean N. Avedissian, Theodoros Xanthos, Athanasios Chalkias, Ernesto d’Aloja, Emanuela Locci, Annette Gilchrist, Walter C. Prozialeck, Nathaniel J. Rhodes, Thomas P. Lodise, Julie C. Fitzgerald, Kevin Downes, Athena F. Zuppa, Marc H. Scheetz

## Abstract

**Introduction:** Vancomycin and piperacillin tazobactam (VAN+TZP) are two of the most commonly utilized antibiotics in the hospital setting and are reported in clinical studies to increase acute kidney injury (AKI). However, no clinical study has demonstrated that synergistic toxicity occurs, only that serum creatinine (SCr) increases with VAN+TZP. The purpose of this study was to assess biologic plausibility by quantifying kidney injury between VAN, TZP, and VAN+TZP treatments using a translational rat model of AKI and rat kidney epithelial cell studies.

**Methods:** (i) Male Sprague-Dawley rats (n=32) received either saline, VAN 150 mg/kg/day intravenously, TZP 1400 mg/kg/day via intraperitoneal injection, or VAN+TZP. Animals were placed in metabolic cages pre-study and on drug dosing days 1-3. Urinary biomarkers and histopathology were analyzed. (ii) Cellular injury of VAN+TZP was assessed in serum-deprived rat kidney cells (NRK-52E) using an alamarBlue® viability assay. Cells were incubated with antibiotics VAN, TZP, cefepime, and gentamicin alone or combined with the same drugs plus VAN 1 mg/mL.

**Results:** In the VAN-treated rats, urinary KIM-1 and clusterin were increased on days 1, 2, and 3 compared to controls (P<0.001). Elevations were seen only after 3 days of treatment with VAN+TZP (P<0.001 KIM-1, P<0.05 clusterin). Histopathology was only elevated in the VAN group when compared to TZP as a control (P=0.04). Results were consistent across biomarkers and histopathology suggesting that adding TZP did not worsen VAN induced AKI and may even be protective. In NRK-52E cells, VAN alone caused moderate cell death with high doses (IC_50_48.76 mg/mL). TZP alone did not cause cellular death under the same conditions. VAN+TZP was not different from VAN alone in NRK-52E cells (P>0.2).

**Conclusions:** VAN+TZP does not cause more kidney injury than VAN alone in a rat model of VIKI or in rat kidney epithelial cells.

## Introduction

Vancomycin (VAN) and piperacillin-tazobactam (TZP) are two of the most commonly utilized antibiotics in hospitalized patients (after the class of fluoroquinolones) (1–6). Meta-analyses (7–11) compiling over 25,000 patients suggest that VAN+TZP increases AKI by an absolute of 9% when considered against VAN + relevant comparators with mean odds ratios ranging from 1.6-3.6 fold higher risk (2–6). This suggested synergistic toxicity is very concerning as experts now recommend avoiding VAN+TZP (12, 13) even though VAN+TZP is a cornerstone of antibiotic therapy in numerous national guidelines. As concerning, relatively few safe alternatives for TZP exist for septic patients. For example, cefepime is associated with dose-dependent neurotoxicity (14–16), fluoroquinolones have numerous black box warnings (17), carbapenems promote broad antibiotic resistance (18), and aminoglycosides are associated with worse AKI (19, 20). Thus, defining the AKI profile for VAN+TZP is imperative as even moderate AKI increases mortality (21–23) and prolongs hospitalization (21, 22, 24).

Despite recommendations to avoid VAN +TZP, biologic plausibility of the increased renal damage has not been established. All studies that have documented an increased risk of AKI with VAN + TZP have solely relied on SCr as a surrogate of renal injury. SCr is neither highly sensitive nor specific for AKI, and a false positive is potential explanation (25). While SCr is easily measured, it is non-specific for kidney injury because transit is defined by secretion and re-absorption (in addition to free filtration) (25). Thus, SCr can be falsely elevated even when the kidney is not injured because of drug competition for renal tubular secretion (26). Unlike VAN, which causes AKI by inducing oxidative stress at the renal proximal tubule, resulting in uromodulin interaction/cast formation (27) and ATN (28), TZP very rarely causes AKI (28, 29). To date, only AIN has been cited with TZP in a few case reports. Given that an absolute increase of AKI with VAN+TZP is very high (i.e. 9% absolute increase based on SCr) (10), it is extremely unlikely that the rare AIN is the driver of toxicity. Furthermore, no clinical study has demonstrated synergistic toxicity occurs because of these mechanisms, only that SCr increases with VAN+TZP.

While serum creatinine (SCr) increases do not always indicate kidney injury, novel urinary biomarkers may be more sensitive and specific for AKI. Additionally, the rat is an excellent model since novel biomarkers and SCr transit properties are conserved between rats and humans (30, 31) in ATN. Further validating the rat-human translational link, urinary biomarkers (e.g. KIM-1 and Clusterin) are qualified for rat (32) and human drug trials (33) by the FDA (i.e. for drug induced AKI). We have previously demonstrated that urinary kidney injury molecule-1 (KIM-1) and osteopontin predict histopathologic damage in a rat model for VIKI (34, 35). The purpose of this study was to assess biologic plausibility by quantifying kidney injury between VAN, TZP and VAN+TZP treatments using cell studies and a translational rat model of AKI.

## Results

### Rat model

In all, 32 rats completed the protocol (Figure 1a, 1b). The constant variance assumption was not upheld for repeated measures ANOVA for KIM-1, clusterin, or osteopontin (P<0.001 for all). Therefore, the mixed model was utilized. Urinary KIM-1 and clusterin differed between the treatment groups as a function of time, whereas osteopontin did not (Table 1a,1b). Changes in these biomarkers over time for each treatment group are graphically displayed in Figure 1c – 1f. In the VAN group, urinary KIM-1 and clusterin were increased on days 1, 2, and 3 when compared to saline (P≤0.001 for all Table 1). For VAN+TZP vs. saline, KIM-1 and clusterin elevations were seen only after 3 days of treatment (P≤0.001 and p=0.04, respectively); however, both were elevated at day 3 compared to baseline values (P≤0.001). Similar findings were obtained with the LOESS model and non-overlapping confidence intervals (Figure 1f). Osteopontin did not increase between treatment groups or treatment days (Figure 1e, P>0.05 for all).

**Figure 1.**
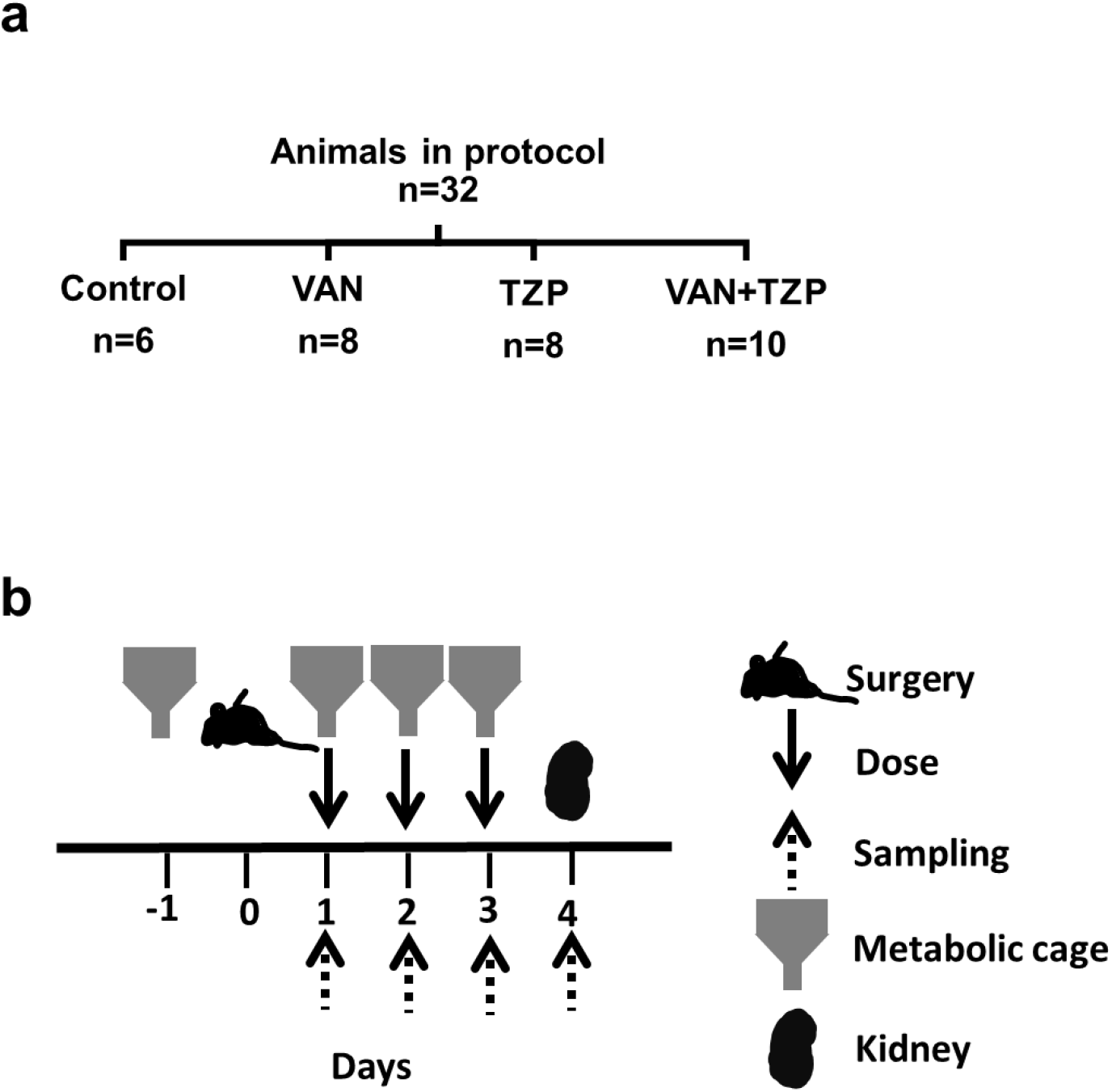
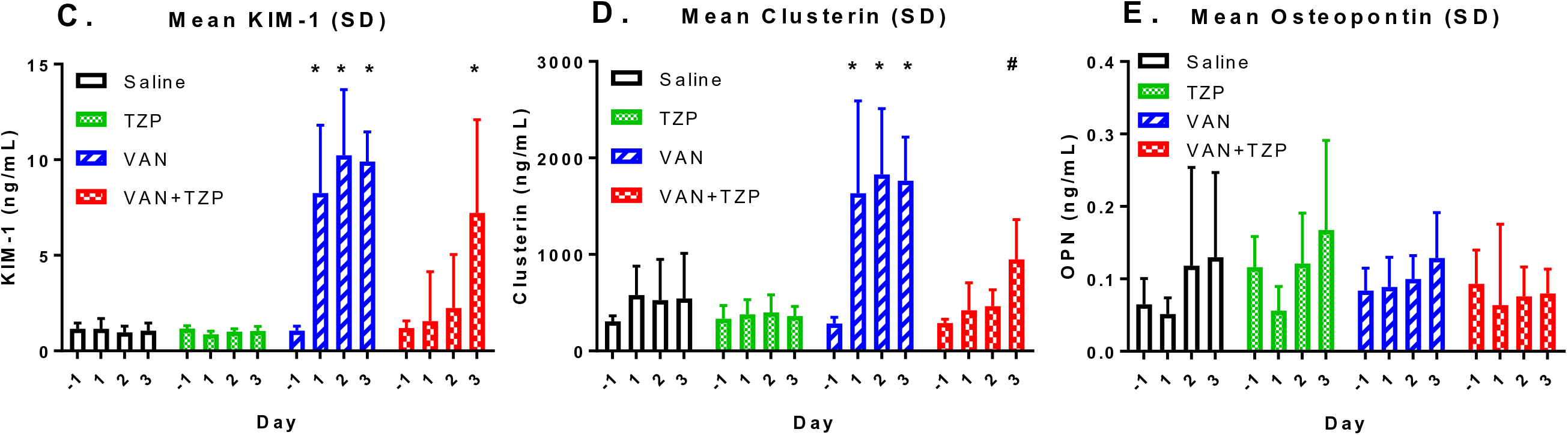
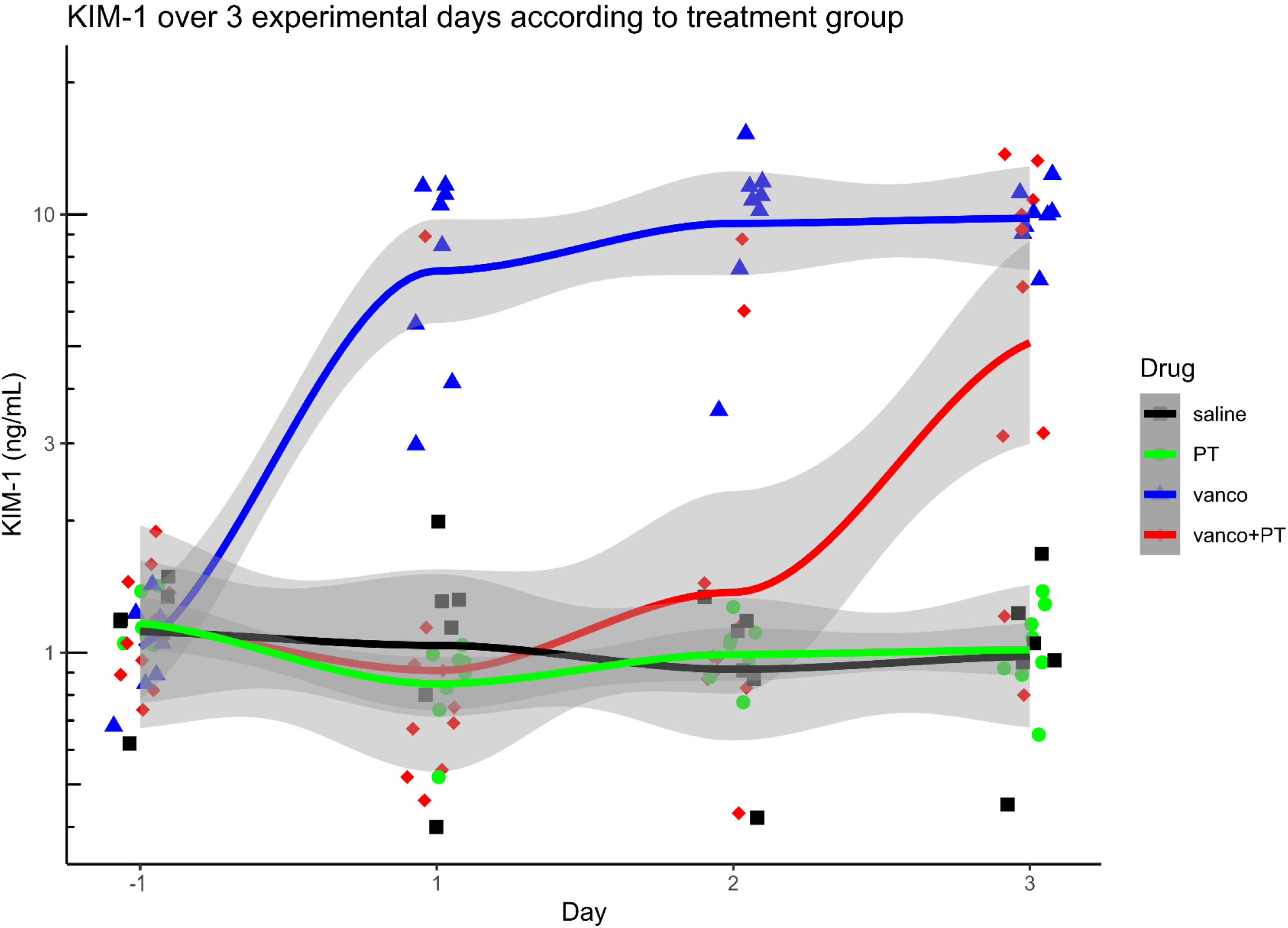
(**a**) Flow chart of animal dosing. (**b**) Timeline of the experiments. (**c**) Urinary KIM-1, and (**d**) urinary clusterin (**e**) urinary Osteopontin levels in saline (n=6), TZP (n=8), VAN (n=8) and VAN +TZP (n=10) - treated animals. Values are expressed as mean ± SD; *p<0.001 vs saline#p<0.05 vs saline, (**f**) Predictive margins with 95% CI of KIM-1 over 3 experimental days according to treatment group KIM-1 = kidney injury molecule-1, CI = confidence interval.

**Table 1.** Contrasts of Marginal Linear Predictions from a Mixed-Effects, Restricted Maximal Likelihood Estimation Regression of Vancomycin (VAN), Piperacillin-tazobactam (TZP), and VAN+TZP on Urinary Biomarkers. Results presented are interactions of drug and day compared to baseline of saline control (Table 1a) and pre-therapy values at day= −1 (Table 1b)

| Drug comparison, Day | KIM-1 |  |  |  | Clusterin |  |  |  | Osteopontin |  |  |  |
| --- | --- | --- | --- | --- | --- | --- | --- | --- | --- | --- | --- | --- |
|  | Contrast | P-value | Lower 95% CI | Upper 95% CI | Contrast | P-value | Lower 95% CI | Upper 95% CI | Contrast | P-value | Lower 95% CI | Upper 95% CI |
| **TZP vs Saline, -1** | 0.020 | 0.987 | -2.301 | 2.341 | 27.012 | 0.897 | -381.849 | 435.873 | 0.051 | 0.172 | -0.022 | 0.125 |
| **TZP vs Saline, 1** | -0.294 | 0.804 | -2.615 | 2.027 | -201.102 | 0.335 | -609.963 | 207.759 | 0.005 | 0.903 | -0.069 | 0.078 |
| **TZP vs Saline, 2** | 0.028 | 0.981 | -2.293 | 2.349 | -128.696 | 0.537 | -537.557 | 280.165 | 0.003 | 0.938 | -0.071 | 0.076 |
| **TZP vs Saline, 3** | -0.013 | 0.991 | -2.334 | 2.308 | -183.387 | 0.379 | -592.248 | 225.474 | 0.038 | 0.318 | -0.036 | 0.111 |
| **VAN vs Saline, -1** | -0.103 | 0.931 | -2.424 | 2.219 | -24.470 | 0.907 | -433.331 | 384.391 | 0.019 | 0.617 | -0.055 | 0.092 |
| **VAN vs Saline, 1** | 7.100 | ≤0.001 | 4.779 | 9.421 | 1054.483 | ≤0.001 | 645.622 | 1463.344 | 0.037 | 0.323 | -0.036 | 0.111 |
| **VAN vs Saline, 2** | 9.260 | ≤0.001 | 6.939 | 11.581 | 1300.684 | ≤0.001 | 891.823 | 1709.545 | -0.018 | 0.625 | -0.092 | 0.055 |
| **VAN vs Saline, 3** | 8.855 | ≤0.001 | 6.534 | 11.176 | 1218.917 | ≤0.001 | 810.056 | 1627.778 | -0.001 | 0.973 | -0.075 | 0.072 |
| **VAN+TZP vs Saline, -1** | 0.040 | 0.972 | -2.179 | 2.259 | -18.003 | 0.928 | -408.949 | 372.943 | 0.028 | 0.435 | -0.042 | 0.098 |
| **VAN+TZP vs Saline, 1** | 0.393 | 0.729 | -1.826 | 2.612 | -158.052 | 0.428 | -548.998 | 232.894 | 0.012 | 0.731 | -0.058 | 0.083 |
| **VAN+TZP vs Saline, 2** | 1.266 | 0.264 | -0.954 | 3.485 | -62.476 | 0.754 | -453.422 | 328.470 | -0.042 | 0.238 | -0.113 | 0.028 |
| **VAN+TZP vs Saline, 3** | 6.160 | ≤0.001 | 3.940 | 8.379 | 402.822 | 0.043 | 11.876 | 793.768 | -0.050 | 0.163 | -0.120 | 0.020 |

| Day Comparison, Drug | KIM-1 |  |  |  | Clusterin |  |  |  | Osteopontin |  |  |  |
| --- | --- | --- | --- | --- | --- | --- | --- | --- | --- | --- | --- | --- |
|  | Contrast | P-value | Lower 95% CI | Upper 95% CI | Contrast | P-value | Lower 95% CI | Upper 95% CI | Contrast | P-value | Lower 95% CI | Upper 95% CI |
| (1 vs. -1) Saline | 0.005 | 0.997 | -2.232 | 2.242 | 272.385 | 0.206 | -150.103 | 694.873 | -0.013 | 0.698 | -0.081 | 0.054 |
| (1 vs. -1) PT | -0.309 | 0.755 | -2.246 | 1.629 | 44.271 | 0.813 | -321.614 | 410.157 | -0.060 | 0.044 | -0.118 | -0.002 |
| (1 vs. -1) V | 7.208 | ≤0.001 | 5.270 | 9.145 | 1351.339 | ≤0.001 | 985.453 | 1717.224 | 0.005 | 0.866 | -0.053 | 0.063 |
| (1 vs. -1) V+PT | 0.358 | 0.686 | -1.375 | 2.091 | 132.336 | 0.428 | -194.922 | 459.594 | -0.029 | 0.275 | -0.081 | 0.023 |
| (2 vs. -1) Saline | -0.182 | 0.874 | -2.419 | 2.056 | 219.802 | 0.308 | -202.687 | 642.290 | 0.053 | 0.120 | -0.014 | 0.121 |
| (2 vs. -1) PT | -0.174 | 0.860 | -2.111 | 1.764 | 64.094 | 0.731 | -301.792 | 429.979 | 0.005 | 0.866 | -0.053 | 0.063 |
| (2 vs. -1) V | 9.181 | ≤0.001 | 7.244 | 11.119 | 1544.956 | ≤0.001 | 1179.071 | 1910.842 | 0.016 | 0.585 | -0.042 | 0.075 |
| (2 vs. -1) V+PT | 1.044 | 0.238 | -0.689 | 2.777 | 175.328 | 0.294 | -151.930 | 502.586 | -0.017 | 0.523 | -0.069 | 0.035 |
| (3 vs. -1) Saline | -0.102 | 0.929 | -2.339 | 2.136 | 237.247 | 0.271 | -185.242 | 659.735 | 0.065 | 0.058 | -0.002 | 0.132 |
| (3 vs. -1) PT | -0.135 | 0.891 | -2.073 | 1.803 | 26.847 | 0.886 | -339.038 | 392.733 | 0.051 | 0.085 | -0.007 | 0.110 |
| (3 vs. -1) V | 8.856 | ≤0.001 | 6.919 | 10.794 | 1480.634 | ≤0.001 | 1114.748 | 1846.519 | 0.045 | 0.130 | -0.013 | 0.103 |
| (3 vs. -1) V+PT | 6.018 | ≤0.001 | 4.285 | 7.751 | 658.071 | ≤0.001 | 330.813 | 985.329 | -0.013 | 0.625 | -0.065 | 0.039 |

Ordinal histopathological scores were increased after treatment with VAN, TZP, and VAN+TZP compared to the controls (Figure 2a). Control animals that received normal saline displayed normal kidney tissue histology (Figure 2b). When TZP was used as the referent group, histopathological scores were significantly elevated in the VAN-treated group (score 2.16, P=0.044), significantly lower in the saline group (score −2.8, P=0.02), and not different in the VAN+TZP group (score 0.29, P=0.76). In the model to predict histopathologic score ≥2, the saline group accurately classified the absence of injury (Table 2). Treatment with VAN was associated with a borderline increased risk of injury (odds ratio 11.67, p=0.058). On the other hand, VAN+TZP treatment was not associated with injury (odds ratio 1.11, P=0.91). The presence of casts was observed in all groups, but the incidence of casts was most prevalent in VAN, either alone or in combination with TZP. A representative cast is visualized in a stained kidney section from a TZP treated rat (Figure 2c). More casts were observed among VAN (8 casts) and VAN+TZP (9 casts) treated groups compared to either saline (2 casts) or TZP (4 casts) treated groups. Tubular dilatation and tubular basophilia were observed in all treatment groups but absent in controls. Notably, tubular basophilia and tubular dilatation were evident in every single kidney section in the VAN group (n=8/8, Figure 2d). However, these findings were less common among the TZP and VAN+TZP treated groups (n=2 and 4/8 and n=2 and 4/10, respectively), suggesting possible nephron-protection. Tubular degeneration was observed only among VAN and VAN+TZP treated groups but not in control and TZP groups. Tubular degeneration was more pronounced in rats treated with VAN alone (Figure 2c) compared to the VAN+TZP-treated group (Figure 2e).

**Figure 2.**
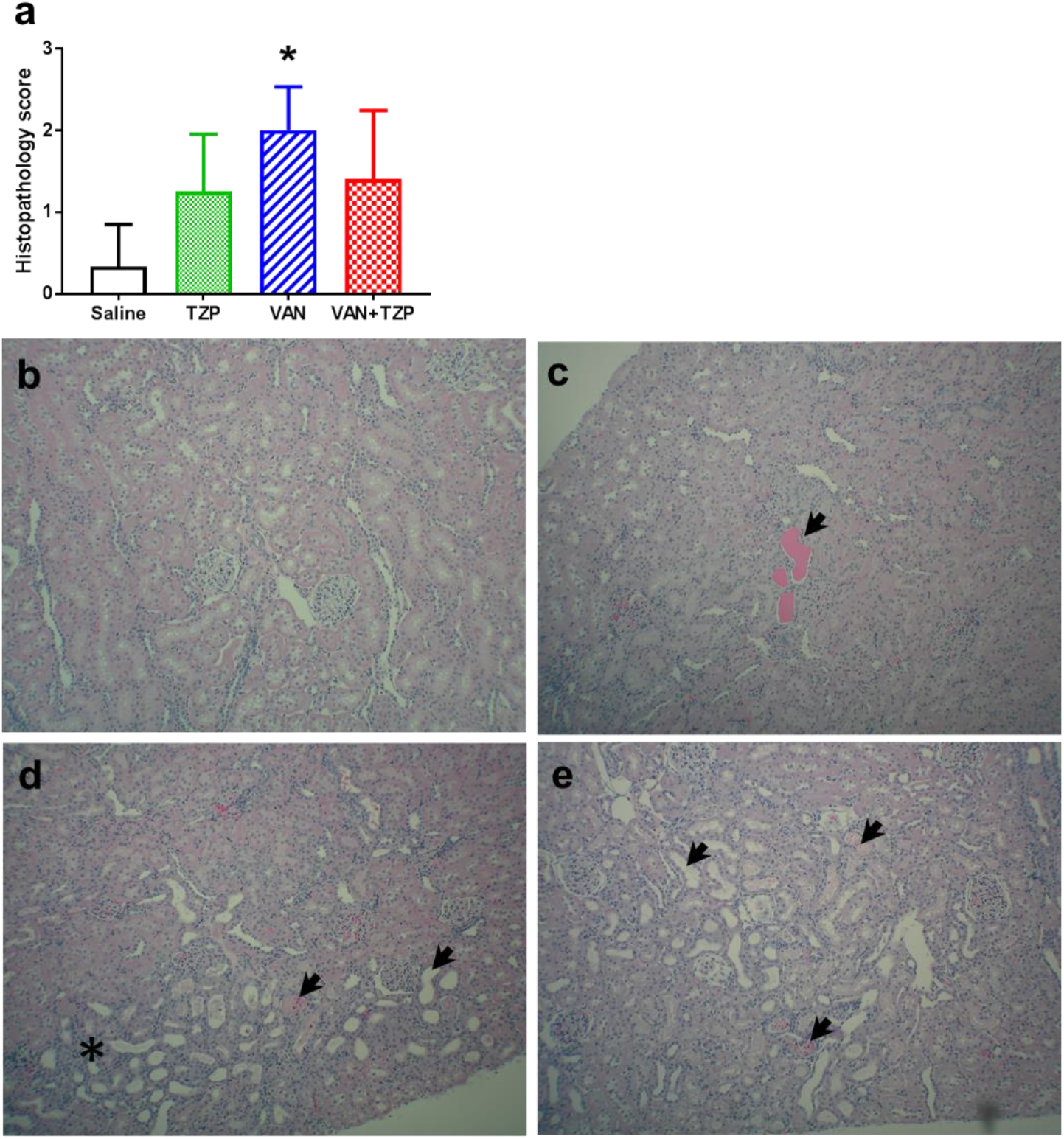
(**a**) Quantitative analysis of histopathology score in kidney tissues of rats. Values are expressed as mean ± SD; *p<0.044 vs saline. Representative photomicrographs of H&E stained rat kidney sections. (**b**) Normal histology of kidney tissue in control rats administered normal saline, (**c**) tubular casts (arrow head, x100) in cortex of rats treated with TZP1400 mg/kg i.p. once daily for 3 days, (**d**) tubular dilatation, tubular degeneration (arrow heads) and tubular basophilia (asterisk, x100) in cortex of rats treated with VAN 150 mg/kg i.v. once daily for 3 days, and (**e**) tubular dilatation, tubular degeneration (arrow heads, x100) in cortex of rats treated with VAN+TZP once daily for 3 days.

**Table 2.** Ordered Logistic regression on Day 3 Histopathology score when comparing to either Saline or TZP

|  | Coefficient | P-Value | Lower 95% CI | Upper 95% CI |
| --- | --- | --- | --- | --- |
| **Groups compared to Saline** |  |  |  |  |
| **TZP** | 2.834 | 0.020 | 0.438 | 5.230 |
| **VAN** | 4.993 | ≤0.001 | 2.287 | 7.699 |
| **VAN+TZP** | 3.123 | 0.010 | 0.753 | 5.494 |
| **Groups compared to TZP** |  |  |  |  |
| **Saline** | -2.834 | 0.020 | -5.230 | -0.438 |
| **VAN** | 2.159 | 0.044 | 0.061 | 4.257 |
| **VAN+TZP** | 0.290 | 0.755 | -1.526 | 2.105 |

### NRK-52E cell experiments

The results of the cellular studies using serum-deprived normal rat kidney NRK-52E epithelial cells are displayed in Figure 3. Treatment with VAN alone produced cell death with an IC_50_ value of 48.76 mg/mL (Figure 3a and 3b). When cells were treated with TZP and cefepime in the absence of VAN, cellular death was not observed (Figure 3a). When VAN was combined with fixed concentrations of TZP or cefepime, the degree of cellular death was not different compared to treatment with VAN alone (Figure 3b, P>0.2). As opposed to VAN treatment, gentamicin produced significant cellular death, producing an IC_50_ value of 11.29 mg/mL. However, when gentamicin was combined with VAN, the IC_50_ of 6.98 mg/mL did not differ compared to gentamicin alone (P>0.2).

**Figure 3.**
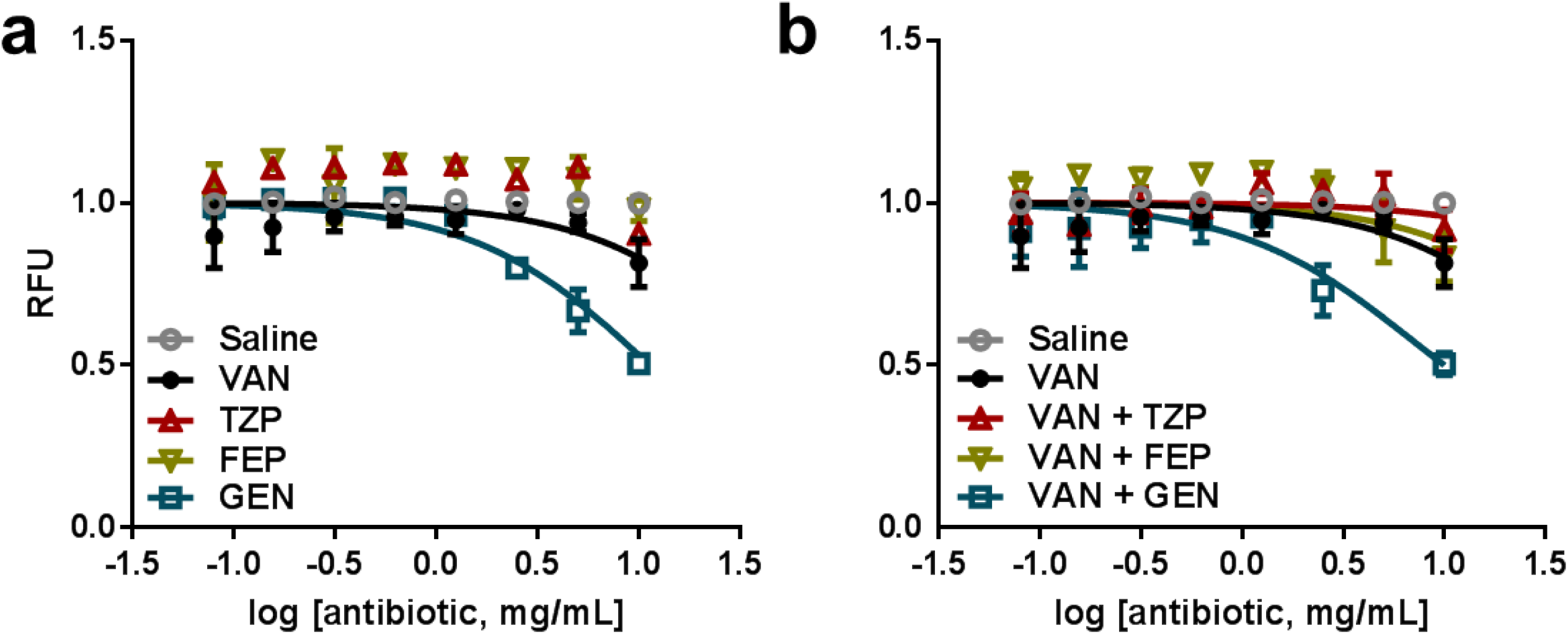
Serum-deprived normal rat kidney NRK-52E epithelial cell viability after treatment with vancomycin (VAN), piperacillin-tazobactam (TZP), cefepime (FEP) and gentamicin (GEN) alone (**a**) or in combination with 1 mg/mL VAN (**b**) for 48 h using alamarBlue® assay. The results presented have been obtained from a single experiment in triplicate but are representative of experiments conducted under different conditions. Values are expressed as mean ± SD; RFU = relative fluorescence units

## Discussion

While a growing number of clinical studies have reported that VAN+TZP is associated with increased rates of AKI as measured by SCr, we demonstrated that VIKI from VAN+TZP is not worse than VAN alone utilizing a translational rat model and a cellular model. In fact, our rat data suggest that TZP may be initially protective against VIKI. The rat data were consistent among relevant urinary biomarkers (i.e. KIM-1 and clusterin) and histopathology. VAN resulted in earlier damage at day 1 compared to VAN+TZP, TZP, or saline. By day 3 for urinary KIM-1 and clusterin, VAN+TZP was similar to VAN alone and elevated compared to TZP alone and saline. On day 3 for histopathology, we observed casts in all rat drug groups, but more casts were seen in the VAN and VAN+TZP groups. Categorical histopathology scores indicated VAN was worse than VAN+TZP and other comparators. The NRK-52E cell line (Figure 3b) findings also demonstrated that VAN+TZP is not worse than VAN alone. Together, these data question the biologic plausibility of VAN+TZP resulting in increased kidney toxicity. To the best of our knowledge, this is the first study investigating the effect of VAN+TZP on kidney injury using translational models focused on histopathology and biomarkers capable of discerning direct toxicity (as opposed to SCr as a surrogate).

Our experimental animal data are consistent with a previous study that demonstrated VIKI in mice but did not identify an increased signal with TZP when using plasma urea as a surrogate for AKI (36). Notably, authors used a VAN 25 mg/day IP for 2 days [which equates to roughly 1000 mg/kg/day or a human allometric scaled dose of ∼80 mg/kg/day (37)] and observed ATN with granular material in the tubular lumen and cast formation. Human VAN doses are ∼60 mg/kg/day even at the upper end of the dosing range (38). This investigation also employed a very low dose of TZP at 100 mg/kg/day IP for 2 days [which is equivalent to 8.2 mg/kg/day human dose (37)]. A standard human dose for a 70 kg patient is ∼ 190 mg/kg/day (39). Thus, it was possible that the very high VAN dose and the very low dose of TZP was the reason for not observing increased toxicity with VAN+TZP. In the present study, we allometrically scaled VAN and TZP to the human dose and did not observe an increase in nephrotoxicity with VAN+TZP (when compared to VAN alone). The results were consistent across the biomarkers and histopathology demonstrating that adding TZP did not worsen VIKI and many even be protective.

We demonstrated that VAN resulted in cell death that fell between positive (i.e. gentamicin) and negative (i.e. cefepime) controls in NRK-52E cells (i.e. rat renal proximal tubule cells). The addition of TZP to VAN did not affect cell death. Similar results were obtained using other cell lines (i.e. HEK-293 and MDCK.2) under different culture conditions (data not shown). Notably, the IC_50_ of VAN from our study is higher than that reported by other groups in kidney cell lines (40–42), but we used clinical grade VAN in our cell studies. The pH of the final solution from clinical grade VAN should be less toxic to cells (43). This removes a variable that could cause cellular death by means other than drug toxicity. Regardless, we did obtain a dose-response-toxicity curve with VAN alone.

Mechanistically, it is well established that VIKI is caused by 1) oxidative stress (28) and uromodulin interaction/cast formation (27) leading to acute tubular necrosis (ATN) (44, 45) and 2) rarely acute interstitial nephritis (AIN) (28). Piperacillin can rarely cause AIN (29). However, no clinical study has studied or demonstrated that synergistic toxicity occurs because of these mechanisms, only that SCr increases. It is notable that TZP on its own is not generally a nephrotoxin (outside of rare cases on AIN). Prospective randomized controlled trial data further support that TZP alone is not overtly damaging to the kidney. For instance, when TZP was compared to meropenem-vaborbactam in 545 patients with urinary tract infections, TZP was reported in adverse events to increase SCr a rate of 0.4% vs. 0% in meropenem-vaborbactam (46). In a separate large trial of 391 patients comparing TZP vs. meropenem for resistant Gram-negative bloodstream infections, TZP only resulted in n=1 non-fatal serious adverse event with increased creatinine vs. n=0 for meropenem (47). At least some large retrospective studies have also failed to find differences in AKI rates (with SCr as a surrogate) (48, 49). Given our findings and those others, the most likely explanation for the reported increase in AKI with VAN+TZP is a false-positive from an imperfect surrogate of kidney injury, i.e. serum creatinine.

Clinical studies generally utilize either the RIFLE or the AKIN definitions to classify acute kidney injury according to SCr (50). SCr is easily measured; though, it is non-specific for kidney injury because transit is defined by secretion and re-absorption (in addition to free filtration) (25). However, in retrospective studies, SCr is often the only reliable variable available to assess a patient’s kidney function. It is possible that competition for secretion or re-absorption can affect SCr in VAN+TZP while kidney function (as measured by glomerular filtration rate) remains unaffected (51) as has been previously suggested (52). A common example is with sulfamethoxazole-trimethoprim, where SCr can be falsely elevated because of xenobiotic competition for renal tubular secretion (26).

We hypothesize that VAN and TZP may be synergistically competing for creatinine. Both piperacillin and tazobactam are substrates for multiple anion transporters (e.g. OAT1 and 3) (53, 54), and anion transporters have been shown to mediate creatinine transit (55, 56) in addition to the more commonly cited pathways such as cation and multidrug and toxin extrusion transporters (e.g. OCT and MATE). Thus, it is possible that basolateral membrane OAT pumps are inhibiting the secretion of creatinine and falsely increasing serum creatinine clinically, though more work is needed to confirm this hypothesis. Further support of this hypothesis is found among patients where antibiotic therapy is discontinued and renal function is re-evaluated. Some serially obtained clinical data support that a false positive SCr elevation may occur with TZP. Jensen et al. found that among critically ill patients who experienced AKI and had antibiotics stopped, those who had received TZP recovered their GFR as calculated by SCr (2.7mL/min/1.73m^2^ per day, P<0.0001) more quickly (57). On the other hand, no GFR recovery was observed among those who had received meropenem (p=0.63) or cefuroxime (p=0.96) (57).

Novel biomarkers (e.g. KIM-1 and clusterin) are qualified for use in preclinical animal (32) and clinical human (33) studies by the FDA to assess for drug induced AKI. These biomarkers are also qualified by the European Medicines Agency (EMA) and the Japanese Pharmaceuticals and Medical Devices Agency (PMDA) for pre-clinical rodent studies (58). The urinary biomarker data from this study are especially interesting since: 1) KIM-1 and clusterin were recently demonstrated as the most sensitive biomarkers for predicting VIKI defined by histopathology (Antimicrob Agents Chemo manuscript invited revision), and 2) multiple samples over time facilitate temporal investigation of toxicity after renal insult. Both of these urinary biomarkers are reasonably specific for tubular toxicity, the location of VIKI (34). KIM-1 is highly sensitive and specific for proximal tubule damage, the locale of VIKI (34). KIM-1 is highly conserved between rats and humans, thus the predictive capacity of rat KIM-1 for VIKI is highly compelling as a pre-clinical model. A human homolog, KIM-1b, is structurally similar except for the cytoplasmic domain (59). Clusterin is present in kidney tubules, is anti-apoptotic, and confers cell protection (60). Since it is not filtered through the glomeruli, elevation should indicate tubular damage (60).

Several limitations exist in these data. These data are from a translational rat model, but urinary biomarker and histopathological data from the rat is a well-accepted surrogate for human pathobiology. By employing multiple biomarkers and histopathology, we assessed for the multiple possible manifestations of VIKI. This study was able to circumvent many of the limitations of current clinical data by nature of being prospective and experimental. Obtaining serial urine over time further strengthens the understanding of the time course of toxicology. Histopathology was elevated for all 3 groups when compared to saline; however, when compared to PT, only vancomycin was elevated. This may reflect the inherent subjectivity of histopathologic assessment as the control group was identified to the pathologist (per standard practice (61)); however, the pathologist was blinded to treatments received. Thus, urinary biomarkers may be a more unbiased assessment of AKI. We are currently limited in that that creatinine values have not yet been analyzed. Creatinine will be assayed from plasma and urine in future studies. Additionally, we have not yet conducted drug assays for VAN or TZP, though these experiments are planned. Finally, while we saw signal for KIM-1 and clusterin in our study, osteopontin did not differ, but it is also not the best biomarker for VIKI (Antimicrob Agents Chemo, manuscript invited revision) and it is not specific for proximal tubule necrosis (62). Finally, we constructed multiple models for our kidney biomarker data. While the constant variance was violated in our repeated measures ANOVA, our mixed model analyses and our LOESS regressions resulted in similar interpretations. VAN resulted in higher KIM-1 and clusterin on days 1 and 2, and VAN+PTZP was not different from VAN by day 3. All models agreed that VAN+PT was not worse and TZP may be protective in initial days.

## Conclusion

VAN+TZP does not cause more kidney injury than VAN alone as evidenced by a translational rat model measuring urinary biomarkers and histopathology. Cellular studies similarly supported that toxicity was not increased by TZP. Novel urinary biomarkers may aid in determining whether higher rates of kidney injury with VAN+TZP are realized in clinical studies.

## Materials and Methods

### Chemicals and reagents

Treatments were clinical grade VAN (Lot#: A000005425, Hospira, Lake Forrest, IL), TZP (Lot#: 7P21TQ, WG Critical Care, LLC, Paramus, NJ and Apotex Corporation, Weston, FL), cefepime (Apotex Corporation, Weston, FL), and gentamicin sulfate USP (Medisca, Plattsburgh, NY) or normal saline (Veterinary 0.9% Sodium Chloride Injection USP, Abbott Laboratories, North Chicago, IL). Other materials were similar to our previous reports (34, 35).

### Experimental design and animals

The animal toxicology study was conducted at Midwestern University in Downers Grove, IL. The study protocol was approved by the Institutional Animal Care and Use Committee (IACUC; Protocol #2295) and conducted in compliance with the National Research Council’s publication, the Guide for the Care and Use of Laboratory Animals, 8^th^ edition (63).

Male Sprague-Dawley rats (290 - 320 g, Envigo, Indianapolis, IN) were randomized to receive VAN 150 mg/kg/day (n=8), TZP 1400 mg/kg/day (n=8), and VAN + TZP at the same doses (n=10) or normal saline (n=6). Surgeries and surgical care were carried out as previously described (35). The VAN 150mg/kg/day dose was chosen based on previous studies (34, 64, 65) and to approximate the human dose (30 mg/kg/day) allometrically scaled for the rat (i.e. 30 mg/kg * 6.2 (rat factor) = 186 mg/kg) (66). The TZP 1400 mg/kg/day was chosen to approximate the human dose (225 mg/kg/day) allometrically scaled for the rat (i.e. 225 mg/kg * 6.2 (rat factor) = 1395 mg/kg). VAN was given IV via the left jugular catheter as this has been shown to result in VIKI in our model, and TZP was given IP to extend residence time. Animals were placed in metabolic cages on day −1 and from day 1 through day 3 (Figure 2a). Rats received the first dose of study drug on day 1. In addition to saline as a control, animals served as their own controls, comparing each study day to pre-therapy (i.e. day −1). Rats were housed in a light and temperature-controlled room for the duration of the study and allowed free access to water and food, including the time in which they resided in metabolic cages (Lab Products Inc., catalogue # 40618-R, Seaford, Delaware).

### Blood and urine collection

Blood samples (0.125 mL) were drawn from a single right-sided internal jugular vein catheter in a sedation-free manner when possible and were prepared as plasma (34, 35). Blood samples were collected over days 1 through 3 with a maximum of 15 samples per animal and plasma stored at −80°C for later analysis (Figure 2a). Urine was collected continuously and aliquoted every 24 hours while the animals resided in the metabolic cages (i.e. 24-hour residence). Urine was centrifuged at 400 × *g* for 5 minutes, and supernatant was stored at −80°C until batch analysis.

### Determination of urinary biomarkers of AKI

Urine samples were analyzed in batch to determine 24-hour concentrations of KIM-1, clusterin, OPN based on standards of known concentrations. Microsphere-based Luminex X-MAP technology was used for the determination of all biomarker concentrations, as previously described (67, 68). Urine samples were aliquoted into 96-well plates supplied with MILLIPLEX® MAP Rat Kidney Toxicity Magnetic Bead Panel 1 (EMD Millipore Corporation, Charles, MO), and analyzed according to the manufacturer’s protocol. MILLIPLEX® Analyst v5.1 Flex software (EMD Millipore Corporation, Charles, MO) was used to calculate standard curves and analyte concentrations.

### Histopathological evaluation

Animals were euthanized via exsanguination through the right atrium while under anesthesia with ketamine/xylazine (100/10 mg/kg, by intraperitoneal injection). Kidneys were then harvested, washed in cold isotonic saline. The left kidney was preserved in 10% formalin solution, and the right kidney was flash frozen in liquid nitrogen for later analysis. Histopathological scoring was performed by IDEXX BioAnalytics (West Sacramento, CA) on paraffin-embedded hematoxylin and eosin-stained kidney sections as previously described (35). In brief the PSTC Standardized Kidney Histopathology Lexicon was utilized with categorical scoring according to grades from 0 to 5 (where the grades for pathological lesions were 0 for no observable pathology, 1 for minimal pathology, 2 for mild pathology, 3 for moderate pathology, 4 for marked pathology, and 5 for severe pathology) (65, 69–73). The composite score for an individual animal was calculated as the highest ordinal score from any kidney site (73).

### Cell culture

Normal rat kidney epithelial cells (NRK-52E, ATCC® CRL1571™) (74) were cultured in Dulbecco’s Modified Eagle’s Medium (DMEM, GenClone, Genesee Scientific) with 5% bovine calf serum in 5% CO2 at 37°C. Penicillin and streptomycin were not included the medium. Cells were either sub-cultured or received fresh growth medium 2-3 times per week. For the cell viability experiment, cells were used between passages 14 to 30.

### Cell viability

NRK-52E cells were plated in 96-well half-volume black plates in DMEM. The next day the media was changed to 10 mM Hepes buffered Hank’s Balanced Salt Solution (HHBS) in the absence of serum. Drugs were dissolved in normal saline and added to the cells. Cefepime was a negative control and gentamicin, a known nephrotoxin (75), a positive control. After 24 h incubation with antibiotics at 37°C, 5% CO_2_, cell viability was assessed using alamarBlue® (5 µl/well, Invitrogen). Cells remained in the same drug-HHBS solution until the end of the experiment. Plates were read at 48 h (24 h after the addition of alamarBlue®) using an Enspire Multimode Plate reader (Perkin Elmer) with Ex530/Em590 filters. Results were analyzed by normalizing the cell metabolism of antibiotic-treated cells to saline controls and expressed as relative fluorescence units (RFU). Drug concentrations were transformed to log base 10. Sigmoidal three-parameter dose response curves were generated after 48 h incubation with antibiotics alone or combined with VAN at a concentration below its IC_50_ (i.e. with VAN 1 mg/mL). The relative toxicity was compared using IC_50_ values i.e., the drug concentration resulting in 50% of maximal reduction in cell viability, obtained using GraphPad Prism version 7.02 (GraphPad Software Inc., La Jolla, CA).

### Statistical analysis

For the rat studies, most statistical analyses were performed using Stata IC 15.1 (except where specifically noted). Decisions to perform analyses with repeated measured ANOVA or mixed models were based on the Breusch-Pagan / Cook-Weisberg test for heteroscedasticity on the ANOVA model. Departure from constant variance at a P<0.05 served as a trigger to use a mixed model. Urinary biomarker elevations were compared across treatment groups using a mixed-effects, restricted maximal likelihood estimation regression, with repeated measures occurring over days as a function of individual rat identification number. Additionally, LOESS models with 95% confidence intervals were created using R version 3.5.1 (76) and the package ggplot2 (77) to circumvent fit assumptions. Mild horizontal perturbation/jitter was applied to the points to enhance visualization. From the mixed model, contrasts of the marginal linear predictions (78) facilitated comparisons of treatment groups vs. controls of saline and pre-treatment values (i.e. day −1). Ordinal logistic regression was used to classify the ordered log-odds of being in a higher histopathologic scoring group according to treatment group. Logistic regression was utilized to determine the odds of having a histopathologic score ≥2 when treatment groups were compared to saline or TZP. All tests were two-tailed, with an *a priori* level of statistical significance set at an alpha of 0.05.

For the *in vivo* studies, graphics were generated and inferential statistics were performed in GraphPad Prism version 7.02 (GraphPad Software Inc., La Jolla, CA) and R 3.4.4. Mean and SD were calculated for triplicate wells. Comparison of IC_50_ values across treatment groups was facilitated by constraining a shared bottom value between 0 and 0.3 and a top of 1 for all groups. The extra sum of squares F-test was utilized to compare independent fits with a global fit whereby a conservative alpha level > 0.2 defined IC_50_ values that did not differ.

